# Early male and female footprints of modern humans across Eurasia and Australasia

**DOI:** 10.1101/2024.05.27.596017

**Authors:** Vicente M. Cabrera

## Abstract

As an alternative to a recent coastal southern route followed by modern humans to colonize Eurasia after an Out of Africa around 60 Kya, and under the premise that the evolutionary rate based coalescent ages slowdown going backwards in time, I propose a new model based on phylogenetic and phylogeographic analyses of uniparental markers in present and past modern human populations across Eurasia and Australasia.

The archaeological record favors a northern route that reached China around 120 kya and then descended latitudinally reaching Southeast Asia and islands around 70-60 kya. These ages coincide with the basal split of the mtDNA macrohaplogroup L3’4* and the origin of the Y-chromosome macrohaplogroup CT* and the subsequent splits in Eurasia of mtDNA haplogroups M and N and Y-chromosome C, D and F clades respectively.

Roughly at the same time modern humans arrived in Australasia other groups retreated southwest returning to Africa carrying with them mtDNA L3 and Y-Chromosome E lineages.

Southeast Asia and Southwest-Central Asia were the subsequent demographic centers for the respective colonization of East and northern Asia and Europe. Across the Ganges-Brahmaputra and the Indus valleys, South Asia was colonized from both migratory centers.

## Introduction

The origin of modern humans in Africa roughly 200 Kya and their exit from this continent approximately 60 kya, with subsequent rapid spread across Eurasia following a southern route, was hypothesized based on mtDNA phylogeny ^1–3^ and phylogeography ^4,5^ . This model has been substantially corroborated by Y-chromosome ^6^ and whole genome ^7,8^ studies. However, this genetic view is in strong contradiction with the model deduced from the human fossil and archaeological records. An origin of modern humans in northwest Africa around 300 Kya has been proposed under archaeological ^9^ and phylogenetic ^10^ grounds. An early out of Africa of modern humans between 90 and 130 kya has strong support in the Near East remains unearthed in Skhul and Qafzeh ^11^. The possible exit of this foray into Eurasia is further supported by the presence in the southern Chinese regions of Zhiren ^12^ and Fuyan ^13^ of modern human fossils dated between 80 and 113 kya. Although the authenticity of these ages was questioned ^14^ they were reaffirmed by a strong reply ^15^. Furthermore, the Chinese ages are congruent with a subsequent southward spread to mainland Southeast Asia and Island Southeast Asia where fossil and archaeological dates are younger but still into this early migration frame. Thus, modern human remains have been recently dated around 80 Kya at the Tam Pà Ling site in northern Laos ^16^, roughly at 68 Kya at Lida Ajer in Sumatra ^17^, perhaps at 65 kya in Callao Cave, in the Philippines ^18^, and deduced from Archaeological evidence around 65

kya at Madjedbebe in northern Australia ^19^. However, based on the coalescent ages obtained for different Eurasian lineages using fossil-calibrated human evolutionary rates, the geneticists usually consider any human dispersal in Eurasia earlier than 60 kya as genetically unsuccessful events which did not contributed to the present-day human genetic pool ^8^, but it should be taken into account that there is still no unanimous acceptance of what mutation rate should be applicable in each case for the estimation of the evolutionary rate ^20,21^ and that a strong time-dependent effect has been detected on the human evolutionary rate ^22^, most probably due to changes in the effective population size ^23^ showing evolutionary rates slower than the used mean for Paleolithic times and faster than the mean for recent historic times ^24^. Under these, more permissive genetic grounds, a more conciliatory model, based on the phylogeny and phylogeography of uniparental genetic markers, has been proposed recently to explain the potential evolutionary and migratory processes of modern humans across the African continent ^24^ continuing outside of Africa and into the Middle East around 120-130 Kya ^25^, which harmonically integrate the uniparental genetic data with the fossil and archaeological records. In this study we extend this model to explain the early spread of modern humans across Eurasia and into Near Oceania and Australia trying to demonstrate that the times and human movements deduced from the mtDNA and Y-chromosome phylogenies and phylogeographies are compatible with the human prehistoric path unearthed by the Archaeology.

## Material and Methods

### The usefulness of the uniparental markers

We are working with uniparental genetic markers, which have been put out of fashion under the assumption that they are just single loci, and that the history of a single genetic locus can differ from that of a population just because of chance or selection. However, the uniparental marker inheritance is different from that of the autosomal polymorphisms. To begin with, the autosomes of any individual are a mixture of the autosomes of their parents which, in turn, are mixtures of the autosomes of their respective parents following, backwards in time, an increasing progression of 2^n^ where n is the number of past generations. In contrast, the uniparental markers have a lineal transmission from father to son or mother to daughter. Due to recombination, genes from autosomes can have quite different genealogical histories but this is not the case for uniparental markers. Indeed, you can follow a lineal transmission for a single autosomal marker, but you cannot establish a consistent genealogical tree with these single variants. On the contrary, comparing two mtDNA or Y-chromosome sequences it is possible to obtain a simple tree branch where on the common trunk are placed the shared variants and, on each branch, the particular polymorphisms of each sequence. This phylogenetic structure can be extended using samples of the same and/or different populations so that you can obtain successive common ancestors for some of them that represent putative real individuals not just gene variants. These genealogical trees are the foundation for studies of male and female migrations from different regions at different times. In addition, the geographic structure observed with uniparental markers allows the identification of wide and small phylogeographic ranges for their clades and subclades, some correlated with specific ethnic groups and even with individual genealogies. A potential weakness of these non-recombining markers is that they only trace ancestors who have left offspring of the same sex. Overall, uniparental markers should continue to be used in demographic studies in combination with autosomal markers.

### Molecular evolutionary rate and number of mutations per lineage

In this study we are dealing with the earliest period of the human expansion across Eurasia. To cope with the time-dependence effect observed for the human mtDNA evolutionary rate ^22^, an empirical algorithm denominated “compound rho” was proposed ^26^ . In it the relative number of ancestors between consecutive coalescent periods in a tree (i/i+1) was used as a measure for population size fluctuations between consecutive coalescent periods, but pending of an automated calculator, the application of this algorithm is laborious. However, it has been observed that the oldest coalescent periods in a tree account for most of the time and genetic variability of the whole tree ^27^. Thus, the mean value of a time-dependent evolutionary rate should be found within those deep periods. An indirect proof in favor of this suggestion is the fact that shallow calibrations on a tree produce underestimated coalescent ages ^28^. Moreover, by comparing ancient mtDNA sequences from different time periods with current sequences from the same haplogroup a significant slowdown of the observed evolutionary rate compared to the expected from a constant rate was found ^23^ . Thus, at the period around 40 kya, the rate between the observed and expected values was 0.44. This result is close to the minimum value (0.50) obtained by dividing the number of ancestors of two consecutive periods (i/i+1). Therefore, when there are not more elaborated calculations, an empirical estimation for the mean value of the evolutionary rate of one species could be a half of the mutation rate in the same species. In the case of the human mtDNA reliable germline mutation rates have been published ^21,29^, giving a consensus average of 1.60 × 10^-8^ (95% CI: 0.30 - 5.34 × 10^-8^) mutations per site per year (msy). However, A pedigree based human mtDNA mutation rate was recently calculated giving a higher value (5.8 × 10^-8^ (95% CI: 3.1 – 10.8 × 10^-8^)) ^30^ than those calculated previously. Curiously, this mutation rate overlaps with the evolutionary rate obtained for recent historical human populations (4.33 × 10^-8^ (95% CI: 3.72 – 4.82 × 10^-8^))^23^. Thus, a mean of these three values was used to estimate the mean evolutionary rate, giving a value of 1.50 × 10^-8^ (95% CI: 0.42 - 9.07 × 10^-8^) msy which was used to calculate all the mtDNA haplogroup coalescence ages. In addition, it was also observed ^23^ that past demographic factors as genetic drift, bottlenecks and founder effects tend to diminish the present-day haplogroup genetic diversity. As a practical approach to correct for these effects, we have used here the most divergent lineages within haplogroups to calculate their coalescent ages.

The most used mutation rates for the Y-chromosome coalescent age estimations are, 1.0 × 10^-9^ (95% CI: 0.3 × 10^-9^ – 2.5 × 10^-9^) msy ^31^ and 8.71 × 10^-10^ (95% CI: 8.03 – 9.43 × 10^-10^) msy ^32^. However, as in the case of the mtDNA, and confirming the time-dependence effect on the Y-chromosome evolutionary rate too, coalescence ages slowdown upwards in time when using ancient DNA calibration ^33^. Thus, following the same rule of thumb as for the mtDNA, a mean Y-chromosome mutation rate was calculated (0.94 × 10^-9^ (95% CI: 0.46 – 1.89 × 10^-9^) msy) and, after halving (0.47 × 10^-9^ (95% CI: 0.21 – 0.95 × 10^-9^) msy), it was used here as a putative mean evolutionary rate to calculate coalescent ages of the Y-chromosome main haplogroups.

### Samples

In this study we used published complete mitogenomes and Y-Chromosome haplotypes obtained from the following databases: NCBI GenBank (www.ncbi.nlm.nih.gov/genbank/, (accessed on 30 June 2023)), Mitomap (www.mitomap.org/MITOMAP, (accessed on 30 June 2023)), Ian Logan 2023 (www.ianlogan.co.uk/sequences_by_group/haplogroup_select.htm,(accesed on 30 June 2023)) and AmtDB (www.http://amtdb.org, (accessed on 30 June 2023)) for mtDNA and the ISOGG database (http://www.isogg.org/tree/, (accessed on 30 June 2023)) for Y-chromosome. In addition, published samples from references listed in the supplementary information were also analyzed.

### Phylogeny

Mitochondrial DNA and Y-chromosome phylogenetic trees were constructed by hand using the median-joining network method ^34^ . Mitochondrial DNA haplogroup nomenclature was according to PhyloTree database Building 17 (http://www.phylotree.org, (accessed on 30 June 2023)^35^. Y-chromosome haplogroup nomenclature was according to ISOGG database (http://www.isogg.org/tree/, (accessed on 30 June 2023)). However, for brevity the Y-chromosome lineages on the text are referred to by the letter of their basic haplogroup and their terminal mutation. Mitochondrial DNA and Y-chromosome coalescence ages were estimated by using statistics rho^36^ and Sigma^37^, and the evolutionary rates proposed above.

### Phylogeography

Based on the fossil remains found in Eurasia, for phylogeographic purposes we divided Eurasia into the following seven geographic regions: 1) Europe, 2) The Middle East-Caucasus, 3) Central Asia (including the Pamir and Tibetan plateaus, the Altai mountains and Mongolia), 4) South Asia (including Afghanistan, Bangladesh, Bhutan, India, Nepal, Pakistan and Sri Lanka) 5) South East Asia (including southern China), 6) Island South East Asia (including ancient Sundaland, the Philippines and Islands of Wallacea), 7) Ancient Sahul (Including New Guinea and Australia), and 8) Near Oceania Islands. As we are dealing with the earliest periods of the modern human spread across Eurasia, we focus on the presence/absence of the mtDNA and Y-chromosome basal lineages in each of the above-mentioned Eurasian regions. To enrich and actualize these analyses we included mtDNA samples reported by YFull MTree (Mitochondrial Tree) and Y-chromosome samples reported by YFull YTree (Ychromosome tree) available at YFull databases https://www.yfull.com, (accessed on 30 June 2023)). Mean haplogroup differences in coalescence ages between regions were calculated by two-tailed t-tests considering that the mean and standard error estimated for haplogroup ages from different samples were normally distributed.

## Results

All databases and the most recent literature about uniparental markers have been screened in search of possible undocumented rare basal lineages. New mtDNA haplogroups and new branches for known mtDNA haplogroups are described for macrohaplogroups M, N, and R, in supplementary figures 1, 2, and 3, respectively.

The most probable geographic origins and their relative ages for lineages within mtDNA macrohaplogroups M, N, and R, are described in supplementary figures 4, 5, and 6, respectively.

The mtDNA sequences used for calculating the coalescent ages of lineages within macrohaplogroups M, N, and R are listed in supplementary tables 1, 2, and 3, respectively.

An actualized phylogenetic tree for the Y chromosome CF-P143 and DE-M145 Eurasian branches and the phylogeography of their respective lineages are depicted in supplementary figure 7.

Coalescent ages for the main Eurasian mtDNA haplogroups in the main geographic areas, and their age comparisons between regions are described in Table1.

**Table 1.** Mitochondrial DNA main haplogroup coalescent ages (kya) in the different regions.

| Regions | West/Central Asia | Southern Asia | Eastern Asia | Southeast Asia | Near Oceania |
| --- | --- | --- | --- | --- | --- |
| **Haplogroup M** |  |  |  |  |  |
| Founder |  | 61.2 | 81.3 | 79.6 | 92.3 |
| 95% CI: |  | 55.8 - 66.7 | 67.9 - 94.5 | 35.0 - 83.8 | 81.2 - 103.3 |
| Radiation |  | 37.7 | 61.1 | 51.3 | 62.9 |
| 95% CI: |  | 32.2 - 43.4 | 43.7 - 78.4 | 44.3 - 58.4 | 41.2 - 84.7 |
| **Haplogroup N** |  |  |  |  |  |
| Founder | 68.5 |  | 53.7 | 66.4 | 68.6 |
| 95% CI: | 56.8 - 80.2 |  | 10.1 - 92.4 | 52.7 - 80.1 | 44.8 - 92.4 |
| Radiation | 39.5 |  | 35.6 | 45.0 | 43.6 |
| 95% CI: | 16.0 - 63.1 |  | 0 - 86.9 | 26.0 - 63.9 | 18.5 - 68.6 |
| **Haplogroup R** |  |  |  |  |  |
| Founder | 88.3 | 89.8 |  | 83.4 | 82.8 |
| 95% CI: | 66.6 - 100.1 | 79.7 - 99.8 |  | 78.2 - 88.7 | 74.6 - 90.8 |
| Radiation | 77.8 | 73.6 |  | 68.0 | 43.3 |
| 95% CI: | 59.5 - 95.7 | 58.9 - 88.4 |  | 58.3 - 77.7 | 4.0 - 82.8 |

Significant comparisons between regions:

Likewise, coalescent ages for the main Eurasian Y-chromosome haplogroups, and their age comparisons between regions are described in Table 2.

**Table 2.**
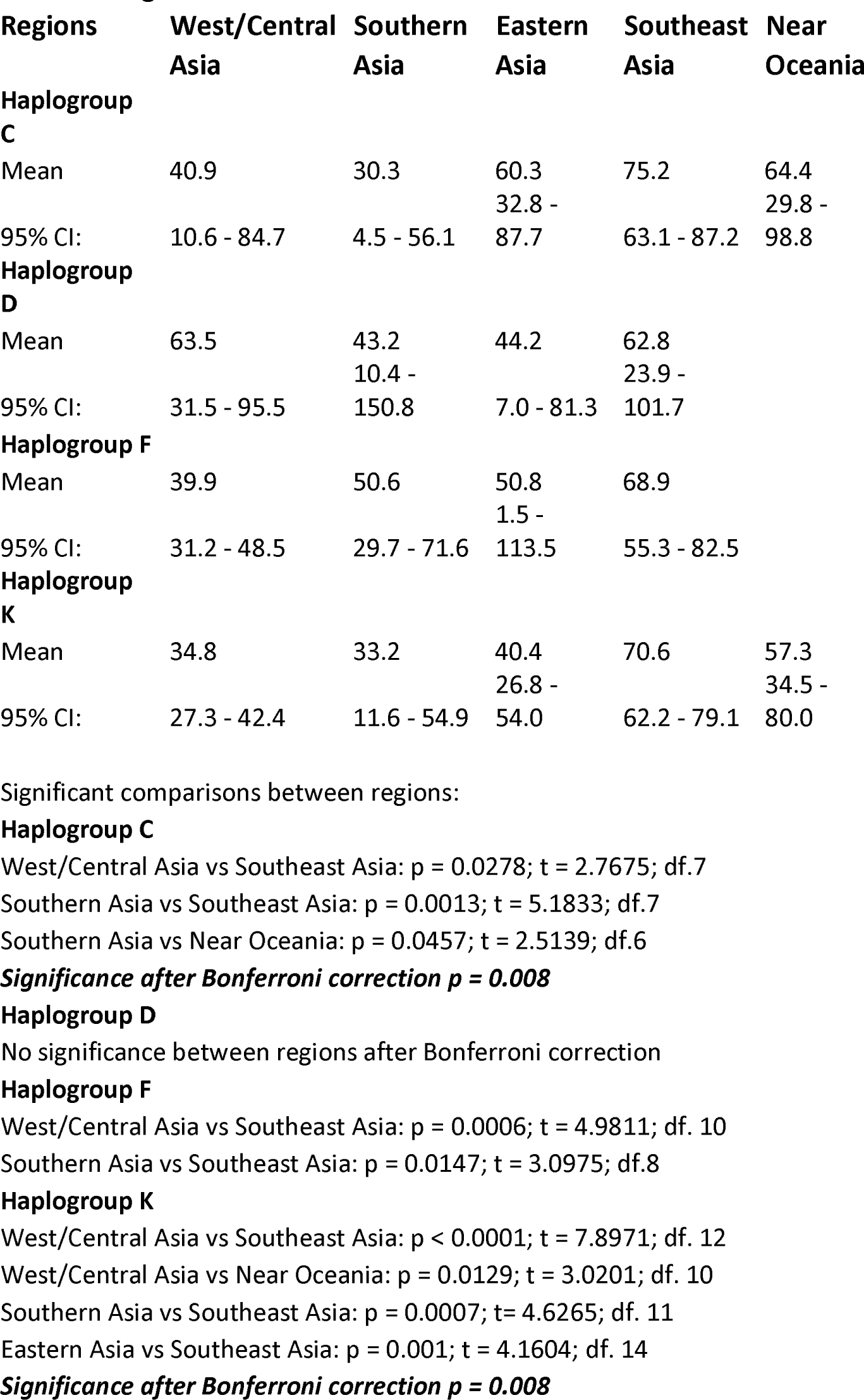
Y-Chromosome main haplogroup coalescent ages (kya) in the different regions.

## Discussion

### The Eurasian fossil record frame

Figure 1 presents the geographic distribution and approximate ages for the main modern human fossil remains unearth across Eurasia.

**Fig 1.**
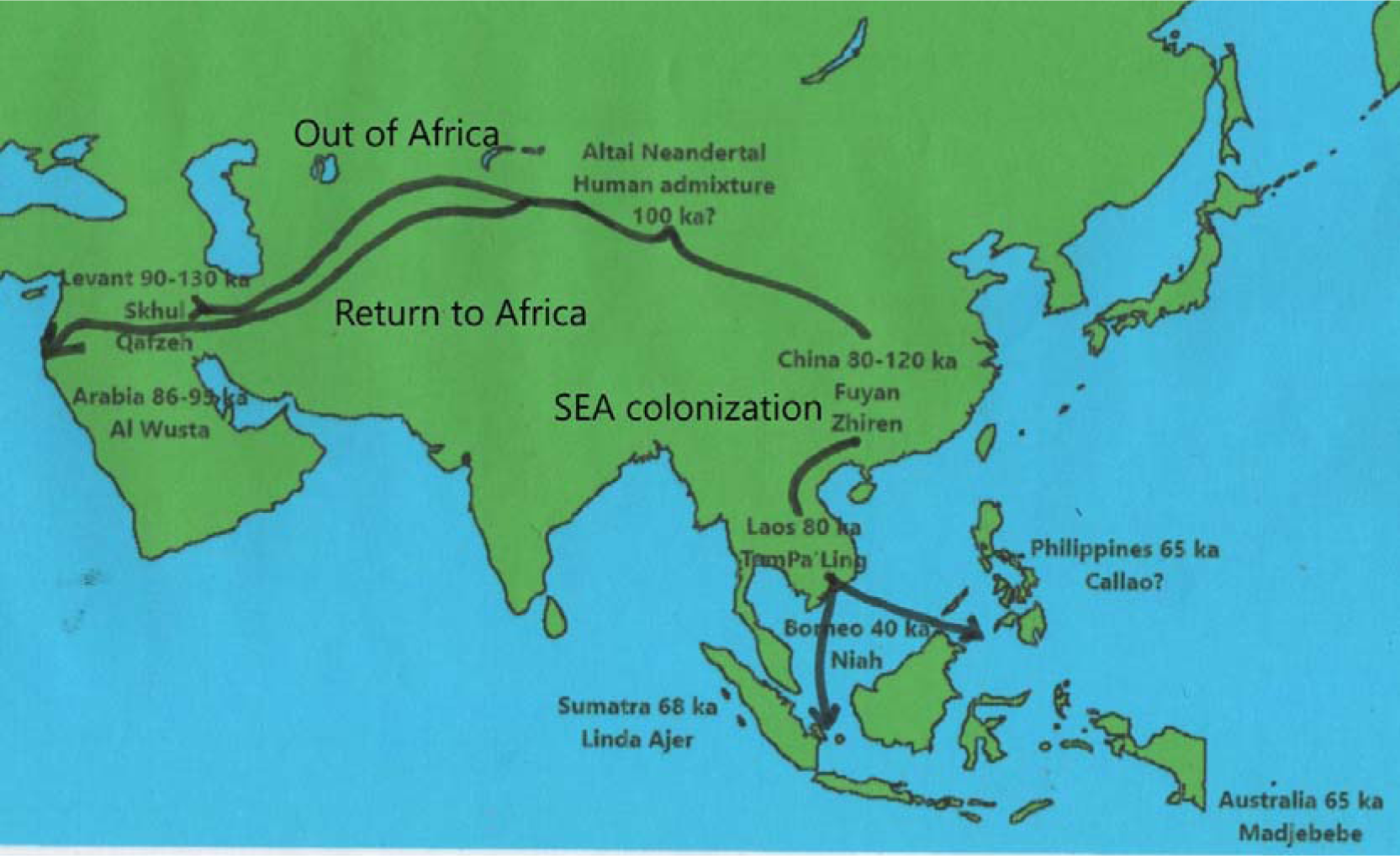
Ages for the oldest human fossil remains unearth across Eurasia. Arrows indicate the first out of Africa northern route and the subsequent return to Africa and colonization of Southeast Asia.

Ages are decreasing longitudinally from the Levant to East Asia and in this region with latitude going southwards to Island Southeast Asia (ISEA). This distribution is more compatible with the hypothesis that modern human followed a northern route to colonize Eurasia after the African exit than the most popular southern coastal route ^38^. In addition, if the out of Africa around 120 kya was a successful exit it would coincide with the climatically favorable Marine Isotope Stage 5e (MIS-5e) facilitating a northward expansion. Furthermore, this northern spread would explain the modern human male introgression on a female Neanderthal genome detected in the Altai Mountains around 100 kya ^39^, and the similarity of the Neanderthal genome segments introgressed into modern human genomes with the approximately 90 ky old Neanderthal genome obtained from the Altai Chagyrskaya Cave specimen ^40^. Furthermore, a very early Neanderthal introgression might have occurred into the ancestors of the 45 ky old Siberian Ust’Ishim specimen around 204.1-95.6 kya ^41^ . How Eurasian modern humans behave at the MIS-5 colder stages d and b is unknown, but there is archaeological information from the coldest stage MIS-4. Around 75 kya Neanderthals went down to the Levant ^42^, a southward retreat that possibly extended to its entire geographic range having a parallel loss of ground by modern humans. Modern humans returned to the Levant around 50 kya ^43,44^ and made inroads into Neanderthal-occupied Europe since that time ^45^. It has been proposed that this modern human secondary spread originated in Africa and recolonized the Levant confronting the Neanderthals. However, at that time it is expected to occur the African uniparental lineages should be basal mtDNA haplogroup L3* and basal Y chromosome haplogroup E* but the European remains supposedly belonging to that African wave had Eurasian lineages as mtDNA N, M and R and Y-chromosome haplogroup C1 (Figure 2). Thus, the molecular evidence favors the hypothesis that the recolonization of the Levant and the first forays into Europe carried out by modern humans originated from a Central Asia core area^46,47^ that could also reach the Near East and northern Africa^48^.

**Fig 2.**
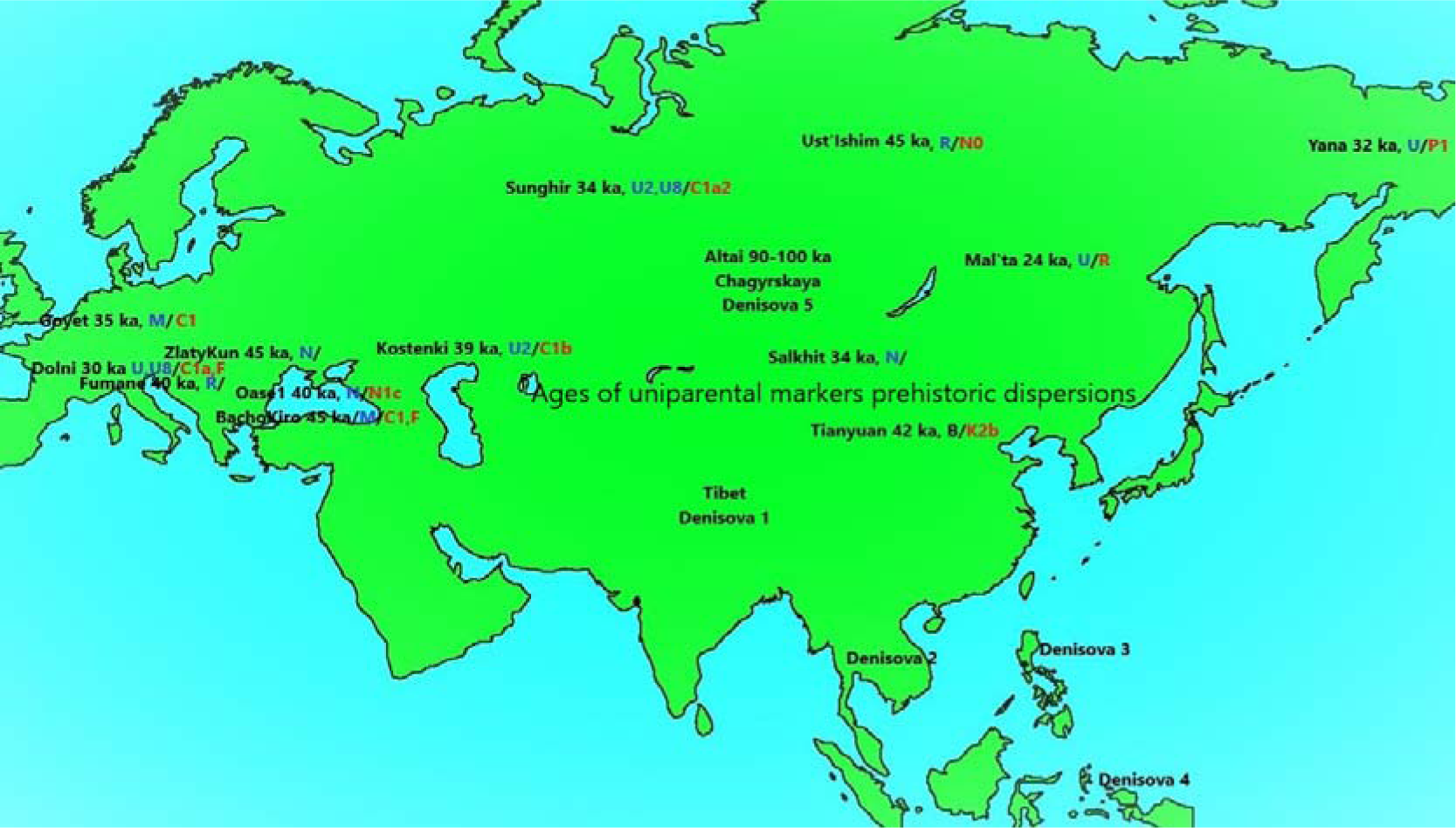
Ancient mtDNA (Black) and Y-Chromosome (Red) lineages obtained from dated modern human remains across Eurasia.

### The ancient DNA Paleolithic window

Due to ancient DNA (aDNA) conservation problems, Paleolithic samples sequenced across Eurasia have a favorable geographic northern bias and a temporal limit around 50 kya (Figure 2). Due to these limitations, and to the deep coalescent ages of the basal uniparental haplogroups, L3* for the mtDNA ^25^ and CT* (CDEF) for the Y-chromosome ^6^, proposed here to be carried by the first out of Africa modern human migrants, it is not unexpected that all mtDNA lineages detected were basal or primitive sequences belonging to macro-haplogroups M or N and N derived lineages belonging to the R macro-haplogroup as U and B (Figure 2). Similarly, Y-chromosome lineages found felt into haplogroups C and F and to the most prominent F derivative, haplogroup K (Figure 2).

However, the Paleolithic geographic distribution of these uniparental lineages contrasts, in some cases, with their current distributions. Outstanding examples are the presence of the present-day western mtDNA haplogroup U in eastern and northeastern Asia as revealed by the analyzed remains from Mal’ta and Yana. On the contrary, today majority eastern mtDNA M lineages were present in Paleolithic European populations as evidenced by the aDNA samples analyzed from Goyet and BachoKiro. A comparable situation occurs for the Y-chromosome results. The currently prominent western haplogroup R was detected in the Siberian remains from Mal’ta ^49^, and basal lineages belonging to haplogroup C, currently dominant in Asia, were detected in the BachoKiro and Goyet European remains ^50,51^ . However, in this case these lineages might persisted and evolved in Europe as attested by the presence of Y-chromosome haplogroup C1a2-V20 haplotypes in Gravettian associated Paleolithic remains from Vestonice (Czechia) and Fournol (France) aged at around 30 kya ^52^ that are still found at low frequencies in present-day European populations. These cases have served to demonstrate that the geographic distribution of the human populations in Paleolithic times could be different to those in present-times ^53^ . In addition, the fact that the specimens analyzed from the BachoKiro cave in Bulgaria^50^ classified in the mtDNA haplogroup N* shared three transitions (4113, 8155, 9456) with the Salkhit (Mongolia) specimen^54^, forming a new branch in the N* tree, provisionally classified as N*3 (Supplementary figure 2), demonstrates the extraordinary migratory capacity of these Paleolithic human groups. Most probably, some of these ancient lineages went extinct. However, more present-day populations must be analyzed before to reach at a definitive conclusion. For example, it was suggested that the Siberian Ust’Ishim specimen did not have modern-day descendants^55^ but later studies demonstrated that he shares 38% of its genome with present-day Siberian and East Asian populations^56^. Furthermore, in the genome of current Tibetan highlanders it was detected an ancient genetic component composed of an admixture of archaic hominins and Ust’Ishim like genomes^57^. In addition, Ust’Ishim and Oase 1 from Romania (Figure 2) share a derived allele at M2308 position within Y-chromosome haplogroup NO with a present-day southeastern Indian Telegu individual^6^ which suggests male genetic continuity and, again, great migratory capacities.

A consequence derived from these aDNA studies was the sequencing of complete genomes of archaic humans as Neanderthals^58^ and Denisovans^59^ which propitiates the discovery of their genetic admixture with modern humans. It is accepted that hybridization with Neanderthals occurred first and in one main pulse^60^, although secondary encounters cannot be ruled out^41,61^. Conversely, the admixture with Denisovans occurred latter and several times across a wide continental range. Denisovans were discovered using only molecular techniques, first from the Denisova cave prehistoric remains at the Altai Mountains in southern Siberia^59^ and later on the Tibetan plateau from Pleistocene remains and sediments^62,63^, but the introgression studies carried out in the genomes of present-day human populations discovered that genetically highly differentiated archaic groups, more or less related to the Altai Denisovans, most probably populated wide additional geographic areas including MSEA, ISEA and even near Oceania^41,64–66^.

### The early out of Africa

Early modern humans left Africa and spread across the Middle East during the humid MIS5e stage around 130 kya^24^. The split of mtDNA haplogroups L3’4 and the origin of Y-chromosome macrohaplogroup CT (Tables 1 and 2) were the molecular uniparental markers of that event. The Sinai Peninsula was one of the most favorable passages for this expansion^67^. The recent confirmation of the existence of basal Y-chromosome haplogroup D2 lineages in western Africa ^68^questioned the most parsimonious exit of only one Y-chromosome composite clade (CT)^6^, favoring instead the exit of three independent lineages (C, D and F). However, the presence of basal D lineages in the Middle East that phylogenetically include the African lineages^69^, the detection of a primitive D1b subclade in the Philippines and possibly in Malaysian Hoabinhian foragers^70^, and the recurrent presence of DE* Y-chromosomes in Tibet^71^ and southern China^72^ are all arguments supporting an Eurasian split of haplogroups D and E that, most probably occurred in Southeast Asia^73^ .

The biparental admixture with Neanderthals in the Caucasus^74^, in the Altai Mountains of southern Siberia^39^, and possibly in Ust’Ishim, western Siberia^41^ strongly points to a subsequent northward spread of these people. However, it is difficult to obtain more direct proofs of this hypothetical northern incursion from any discipline. At molecular level, there is no present or past evidence of early phylogenetic branching of those uniparental markers, probably due to the low population size of those human groups^75^. On the archaeological side, it has been documented that at Middle Paleolithic times early modern and archaic humans used indistinguishable lithic industries^76^, and the mixed features found in numerous remains of that epoch makes difficult its morphological classification.

The lack of basal uniparental lineages in the current populations of Central Asia indicates that those pioneers did not survive to the present-day.

### The first return to Africa

Climatic conditions worsens since MIS5d stage at around 110 kya. Colder conditions could oblige humans to retreat from their northernmost colonized borders, pushing back further southern groups. Avoiding mountain barriers such as the Pamirs and the Himalayas, those migratory movements took some groups of modern humans to Southeast Asia while others returned to the African Continent. This backflow to Africa was first suggested from the Y-chromosome phylogeny and phylogeography ^77^. Later, it was proposed that mtDNA haplogroup L3 could be the female counterpart of that return^25^. Consequently, we have dated this retro-migration around the radiation ages of mtDNA haplogroup L3 and Y-chromosome haplogroup E (Tables 1 and 2). This gene flow from Eurasia to Africa could explain the small Neanderthal component detected in African populations^78^. In addition, Neanderthals could play a competitive role in this modern human retreat to Africa as there is archaeological evidence that modern humans abandoned the Levantine region around 80 kya being replaced by Neanderthals^79^.

Although signaling later migrations, it is worth mentioning that E1b1b-M215 Y-chromosomes were present in the Middle East at least since the Mesolithic Natufian period^80^, and that more derived branches of this haplogroup have been documented in Europe and West Asia. If these branches were the result of sub-Saharan Africa gene flow it should be expected that at least mtDNA haplogroup L3 lineages would also appear in these regions, but this is not the case. Thus, most probably, the accompanied maternal lineages of that spread were of Eurasian origin. As haplogroup E Y-chromosomes represent a key component of the African paternal gene pool, its presence in Europe might explain the fact that genetic distances between Europeans and Africans are lower than those of the later with East Asians or Oceanians^81^. If this hypothesis were correct, a bias due to gender should also be detected in the analysis^82^

### The first expansion in Southeast Asia

Despite the time molecular eclipse commented above, colonizer groups had to migrate at a good pace under adverse conditions and grow fast in favorable places, to form isolated communities where common uniparental lineages diversified independently as found at continental and subcontinental scales. It is deduced from phylogenetic and phylogeographic information that Southeast Asia, including southern China, was one of the regions where the founder lineages first arrived and expanded. For instance, here, after the classification of 232 previously undetermined complete mtDNA sequences, it was possible to construct new basal haplogroup trees or new basal branches of known haplogroups. From these, nineteen belonged to macro-haplogroup M (SFig.1) being 12 (63%) of Southeast Asia adscription and 7 (37%) of South Asia provenance. Only 7 new clades were constructed within macro-haplogroup N (Sfig2), 3 (43%) originated in Southeast Asia and the rest included West Eurasian and Near Oceania samples but none was from South Asia. Eleven new clusters were found in macro-haplogroup R, 6 (55%) had Southeast Asian origin and 2 (18%) were from South Asia (Sfig3). Statistical comparisons between main geographic regions (Table 1) showed that for mtDNA macro-haplogroup M, the founder and radiation coalescent ages of Southeast Asia and Near Oceania M haplogroups are significantly older than in other regions. The fact that human mtDNA M lineages in India have significantly younger ages than those in East Asia, Southeast Asia, and near Oceania, was previously used as evidence against the southern route hypothesis for the colonization of Eurasia and the evidence of an earlier ancestral center of radiation in Southeast Asia^83^. Likewise, the lack of basal mtDNA macro-haplogroup N lineages in India and its presence in Southeast Asia and Near Oceania was used as an argument supporting the existence of a northern route for the colonization of Eurasia^84^. Furthermore, coeval independent dispersals of mtDNA R haplogroups in West Asia and Near Oceania also pointed to the existence of a halfway core-area of expansion in Southeast Asia^85^.

Overlapping phylogeography and mean coalescence ages for the main basal Y-chromosome Eurasian lineages (C, D, F, K) are also in support of an early arrival and early expansion of modern human males in Southeast Asia and Near Oceania (Table 2). The existence of an early center of Y-chromosome expansion in Southeast Asia was first proposed from the result of a high-resolution phylogenetic and phylogeographic analysis of haplogroup K-M526^86^, and afterward for the exclusive presence in this region of confirmed basal F* lineages^6^. In contrast, the reanalysis of Indian putatively basal F* male lineages demonstrated that they were basal members of haplogroup H, a derived branch of macro-haplogroup F^87^. Once more, these results are against the southern route hypothesis throughout the Indian subcontinent.

Thus, all subsequent migratory waves had Southeast Asia as their demographic center of expansion.

### The earliest Asian expansion

The first great split of the early modern human group that colonized Eurasia occurred in Central Asia, and while one subgroup returned to the Middle East and Africa, another group advanced eastwards migrating along the northern slopes of the Himalayas reaching southern China, the Indochina peninsula and Sundaland (Figure 2). After this, the first detectable uniparental expansion from that region is marked by the deep phylogenetic divergence and vast but fragmented geographic distribution, with prominent pockets in the Andaman Islands, Tibet, and Japan, of the Y-chromosome haplogroup D-MCTS3946^71^. Waiting for more accurate and unbiased Y-chromosome sequencing analysis that definitively resolve the identity of the D*(xM174) lineages detected in Asia^88,89^, the most ancestral Y-chromosome D clade in Southeast Asia was found in the Philippines (D1b-L1378), thus, a southeastern region, including the Philippines, may be considered the radiation center for that early migratory wave that had to be close in time to the one that, spreading eastwards, colonized Australasia. D-M174 was detected in a Malaysian Hoabinhian hunter gather remain^70^. From the D-M174 ancestral node two sister branches diverged around 79.8 kya given place to the ancestors of Andamanese and Japanese lineages D1a2b-Y34637 and D1a2a-M64 respectively^90^. This favors the existence of a coastal route with ample latitudinal range. Barely after, a third branch, D-Y15407, gave rise to the two Tibetan clades D1a1a-M15 and D1a1b-P99. Additionally, the detection of an ancestral Y-chromosome Hg P-295* lineage in an historical Andamanese remain^91^, whose phylogenetic counterparts have been only detected in Malaysia and the Philippines (SFig 7), strongly reinforces the hypothesis that the Andaman Archipelago was colonized by a demic westward spread from Sundaland. It is difficult to assign a unique maternal counterpart to the Y-chromosome haplogroup D in the Andaman and Japan as the most prominent mtDNA lineages are different in each region. Secondary branches of Indian/Indochina mtDNA haplogroups M31 and M32 are the maternal representatives in the Onge of Andaman^92^, while secondary branches of mtDNA haplogroups N9b and M7a are the prominent clades in ancient Jomon and present-day Ainu Japanese^93^. However, a paradoxically widespread mtDNA M clade, haplogroup M13, with peaks in frequency and diversity in Tibet and Japan, has been signaled as a possible mtDNA counterpart of the Y-chromosome haplogroup D expansion^94^. In addition, it must be mentioned that a rare mtDNA M lineage (SRG059) detected in West Papua New Guinea^95^ has a very conservative transition, T14440C, that is a diagnostic mutation defining the Indian Indochinese-Onge haplogroup M31. On the other hand, a Holocene hunter-gather sample from the Wallacean Sulawesi Island also showed a deeply divergent mtDNA M lineage^96^ which shares G15777A transition with the common trunk of the new Indochinese proposed here clade M*2 (SFig 1). Both cases are compatible with the existence of a primitive center of radiation in Southeast Asia-Sunda shelf. A prolonged period of isolation and genetic drift followed by independent migrations in each region could explain these results. Note that, although the coalescent age of Y-chromosome haplogroup D is very old, its expansion ages in each region (about 32 kya in Japan and 10 kya in Andaman), are much more recent. The case of Tibet deserves special comment. It has been proposed that a Tibetan specific mtDNA basal haplogroup M lineage (M62) could be the maternal counterpart of the Y-Chromosome haplogroup D^97^. In principle, M62 and the Southeast Asian mtDNA haplogroup M68 conformed a composite haplogroup (M62’68) defined by transitions at 150, 4561, and 7664 positions^35^. However, after the addition of new sequences to the phylogeny of both groups (SFig 1), the M68a branch lacks transition 7664 and a more parsimonious alternative could be to join M62 and M25 (M25’65), as both share transitions at 150, 3511, and 13708 positions (SFig 1). Haplogroup M25 is considered an ancient autochthonous Melanesian lineage^98^, that expanded in the Solomon Islands 10.3 kya (CI: 7.4-13.3 kya). This link suggests a potential gene flow between Tibet and Melanesia. Curiously, this is not the unique case. A rare mtDNA lineage belonged to macrohaplogroup N has been recently reported in Papuans from New Guinea^95^. This lineage, aggregated to an also rare Nepalese sequence, could conform a new haplogroup (N*1 in SFig 2) having transitions at 12681 and 15262 positions as diagnostic mutations. Furthermore, the fact that the mtDNA haplogroup N11a, sister branch of the specifically Philippine N11b clade, is found in Tibet and surrounding regions, and the presence of the rare Y-chromosome P1a-M65 haplogroup in the Philippines, Melanesia, Nepal^99^ and in the Andaman Islands (Sfig 7), all point to an old genetic relationship among these geographically distant regions. In the first mtDNA studies of the Indigenous Andamanese, it was proposed that they represented the direct descendants from the first humans that migrated out of Africa ^100^. However, that hypothesis was soon rejected in favor of a Paleolithic origin from the Indian subcontinent based on more complete mtDNA studies^92,101,102^ or a Southeast Asia origin based on genome studies that showed closest affinities of Andaman Onge people with Malaysian negrito^99^ and ancient Hoabinhian hunter gathers from Laos and Malaysia^70^. Interestingly, in the last study a genome component of Hoabinhian ancestry was detected in ancient Japanese Jomon^70^. Linguistic studies on the nearly extinct Kussunda people from Nepan and Onge from the Andaman Islands demonstrated their affiliation to the Melanesian Indo-Pacific linguistic family, and the possibility that these people were the remnants of the Australo-Melanesian primitive settlers was suggested^103^, but, just the contrary, that they resulted from a Paleolithic westward expansion of the Australo-Melanesian ancestor is equally possible. In sum, the analyzed genetic data are compatible with early human expansions from Southeast Asia/Sunda shelf toward the East and the West as proposed here.

### The early Near Oceania colonization

One of the first arguments questioning the classical southern route hypothesis^104,105^, was the detection in Island Melanesia of very ancient and divergent mtDNA macro-haplogroup M lineages (M27, M28, M29, Q)^106^ and P lineages, belonging to macro-haplogroup R^107^, in that area. When these studies were extended to Australia^108–110^, it was evident that the female colonizers of this far away from Africa Pacific area carried basal mtDNA lineages that directly sprout from the root of the three Eurasian mtDNA lineages M, N and R. Another unexpected observation was the deep genetic isolation between the Melanesian and Australian regions. Y-chromosome studies replicated fairly well the mtDNA results. It was demonstrated since the beginning the profound divergence of the Y-chromosome lineages in Island Melanesia^111^, and the independent histories of Y-chromosome in Melanesia and Australia^112^. These results were confirmed subsequently using high resolution typing and high coverage methods in Australian^109,113^ and Melanesian populations^114,115^.

One of the first questions that aroused interest on the settlement of this area was to know whether it was conducted in one or several waves and what was or were the route/s followed by those early colonizers. The phylogeography of the uniparental markers give some clues to answer these questions. Focusing first on mtDNA, the oldest haplogroup M lineages are found in Island Melanesia, and the dominant M cluster in New Guinea, which could be its most probable conduit to Island Melanesia, is haplogroup Q, an early branch of the Island Melanesia haplogroup M29^106^. Thus, the spread of haplogroup Q in New Guinea may be better explained as a westward introduction from Island Melanesia. One way to explain this surprising result is to suppose that those pioneers who first reached and grew up on the Melanesia Island region arrived there navigating along the northern coast of New Guinea, without permanently penetrating the island. If this hypothesis is accepted, the most probable route followed by those seafarers was the northern route across Sulawesi and Maluku, carrying basal mtDNA M* lineages that, under favorable conditions, radiated first in Island Melanesia. Curiously, a recent genomic analysis of a middle Holocene hunter-gatherer from the Leang Panninge cave in Sulawesi detected a basal mtDNA haplogroup M* in this individual^96^, that shares the conservative transition 6374 with the Island Melanesia M28 clade. Focusing on Australia, it has been demonstrated that this Continent showed a strong Indigenous mtDNA structure^110,116–118^, and that a primary potential center of expansion could be situated in northeast Queensland^117^. Precisely, the autochthonous mtDNA haplogroup M42 is particularly frequent and diverse in this territory^110^ and, congruently, its two main lineages (M42a and M42c) were documented in the Australian Barrineans^119^, so it does not seem unreasonable to propose that the primary mtDNA M* expansion that occurred in Island Melanesia also extended southwards colonizing the northeast of Australia. Is there a potential itinerary overlap signaled by Y-Chromosome markers? Y-chromosome haplogroup SM-PR2099 (SFig.7) seems to be the best candidate as its two main branches, M and S (SFig. 7), are documented in Island Melanesia being M-P256, well represented in New Guinea and scarce and more derived in Australia, an accurate reflection of the geographic distribution of mtDNA haplogroup Q, and S1-B255, more abundant and widespread in Australia, the best counterpart for the Australian mtDNA haplogroup M42. However, this potential northern route does not explain the phylogeography for other Australasian parental lineages. For instance, mtDNA macro-haplogroup N is represented in Australia by three relatively frequent lineages (N13, O and S)^108,110,116,117,120^ that have a clear northwest geographic distribution pointing to this area as a potential point of arrival at the continent^85,117^. However, except for sporadic appearances of N13^95,114^, these lineages are absent in New Guinea. However, a rare N*1 lineage (SFig 2) has been recently detected in New Guinea^95^ with possible phylogenetic affinities with Nepalese lineages (Sfig 2). On the contrary, haplogroup P, a basal branch of macro-haplogroup R in the near Pacific, could have shared the same point of entrance but with a somehow different distribution. Several autochthonous branches of P radiated early in Australia and slightly later in the New Guinea highlands^121^, suggesting that the colonizers who arrived on the western coast of Sahul penetrated inland and branching out, some advanced toward the north and others toward the south. The geographic distribution of other derived mtDNA R lineages with small frequencies as R12 and R14 in Australia and New Guinea respectively, could have followed this second route as well. The Y-chromosome companion of this western side settlement could be derived lineages of haplogroup C1b-F1370 (SFig 7) and haplogroup C1b2-B477 (SFig 7). However, from the uniparental information gathered at present, it is difficult to trace the precise route followed by these colonizers from Sundaland to west Sahul. For example, mtDNA haplogroup N lineages present in Nusa Tenggara (N21 and N22) are younger than the Australian N lineages. The only region in the area that brings together ancestral lineages for the three mitochondrial macro-haplogroups, M (M80), N (N11), R (P9, 10) is the Philippine Archipelago. Its closest mtDNA affinity with Timor-Leste^122^ could be in favor of a southern route throughout Nusa Tenggara for the western settlement of Sahul. Likewise, the presence on the Philippines of the Y-chromosome basal lineages C2-M217 and P-PF5870 that are, respectively, sister branches of lineages C1-F3393 and SM-PR2099 involved in the settlement of Sahul (Sfig 7), points to a main role of the Philippines as a main step on the colonization of Near Oceania. However, another possibility could be that the Philippines, still today, preserve genetic vestiges of the first colonization of modern humans better than other islands in Southeast Asia^123,124^. Thus, against the best fit model based on genomic data that proposed a sole founding wave of modern humans into the Sahul^125^, phylogeography of uniparental markers strongly points to the existence of two waves, although the radiating ages of the lineages involved do not allow to separate them in time (Fig. 3).

**Fig. 3.**
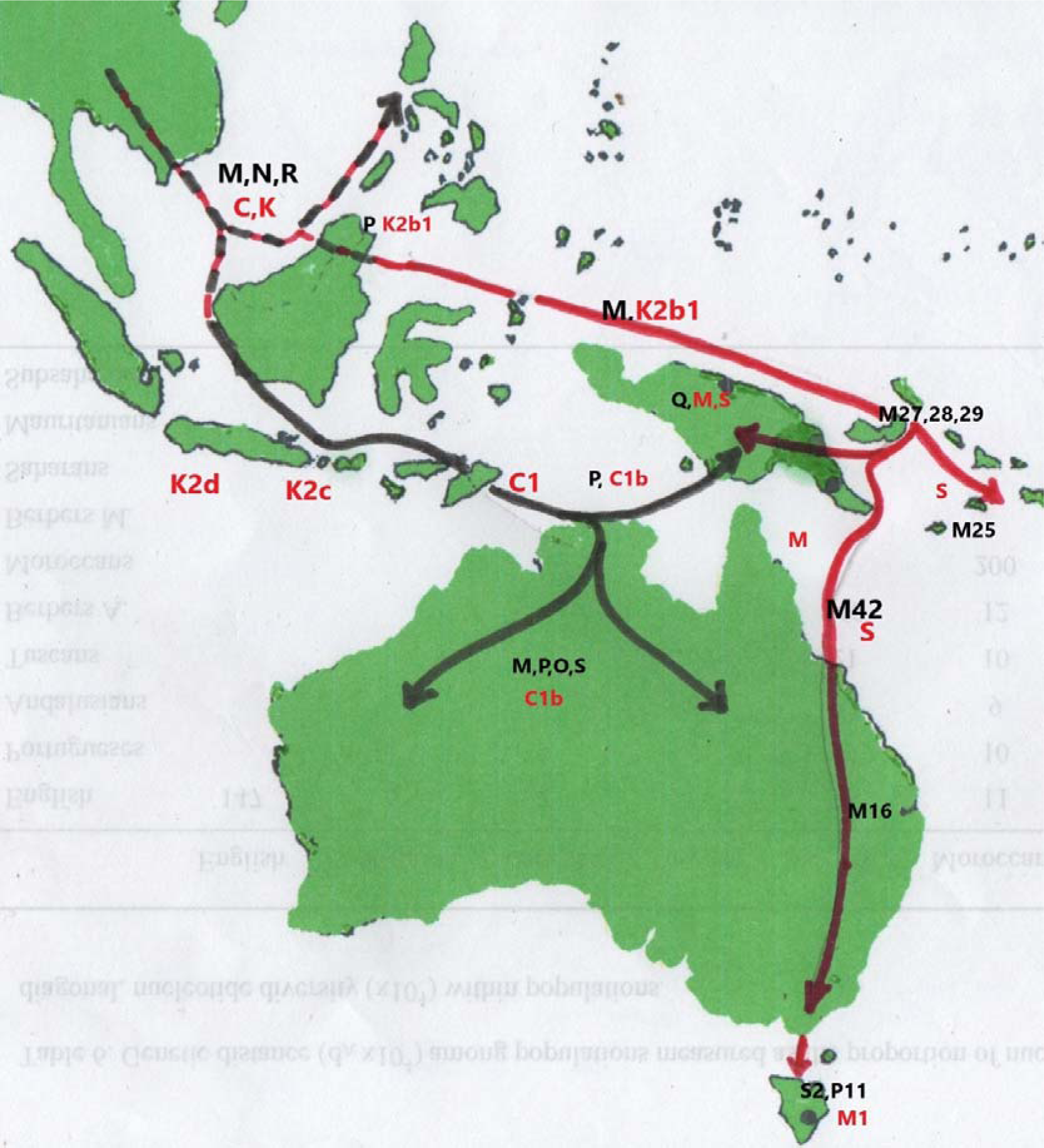
Australasia and Near Oceania colonization following two putative northern (red) and southern (black) routes.

### The first colonization of India

According to the southern coastal route hypothesis, India is considered an obligatory step in the colonization of Eurasia and Australasia by the modern humans who left Africa. However, as previously explained, mtDNA genetic data obtained from Indian populations do not support this hypothesis. First, there are not autochthonous mtDNA macro-haplogroup N(xR) lineages in India and the N(xR) branches present in it have their ancestral roots outside of this subcontinent^84^. It was for this reason that a second northern route, carrying mtDNA macro-haplogroup N lineages to the East, bypassing India, was proposed^94,126^. Second, the Indian macro-haplogroup M lineages, although arising directly from the ancestral M node, have foundation and expansion coalescent ages significantly younger than their counterparts in Southeast Asia and Australasia (Stable 1), while macro-haplogroup M is practically absent from West Eurasia^83^. Third, the majority of the macro-haplogroup R lineages in India shared deep roots with those present in western geographic areas but do not with those to the eastern side^85^.These mtDNA data were reconciled in a hypothesis which proposed that prehistoric modern humans colonized India following the two natural corridors marked, respectively, by the Indus and Ganges rivers at the northwest and northeast sides of South Asia. Macro-haplogroup M lineages entered the subcontinent from the East and those of macro-haplogroup R mainly from the West^83^.

Although without a physical genetic linkage, one would expect a follow you follow me behavior for sex-linked markers. In consequence, a Y-chromosome correlated overlap to the mtDNA Indian phylogeography might be found. Y-chromosome lineages considered indigenous to India belong to haplogroups C, H, L and R^127,128^. As we are dealing with the most primitive settlers, we can discard the haplogroup L1a-M2481 Indian branches as the most ancestral lineages of this haplogroup have been detected in western Pakistan^129^, and the same occurs for haplogrup R1a-M417 for which the roots of the Indian lineages were found in the vicinity of present-day Iran^130^. This left us three possible autochthonous Y-chromosome Indian haplogroups belonging to C1b-F1370, H1a-M69, and R2-M479 clades. The haplogrup C Indian branch, C1b1a1a-M356 has an ancient coalescence age of around 54 kya in India (Table 2) but the ancestral branches for this haplogroup were found in Southeast Asia and Southeast Asian Islands^73^ which clearly points to an entrance into the Indian subcontinent from the East likewise was found for mtDNA haplogroup M lineages^83^. Y-chromosome haplogroup R2-L722 has a more restricted geographic distribution, around and within the Indian subcontinent. This haplogroup arose from the basal R node, defined by the M207 SNP, for which a Central Asia origin was postulated^131^. This origin has been reinforced by the presence of this ancestral R lineage in the Mal’ta 1 Paleolithic specimen unearthed in southern Siberia, west of Lake Baikal^49^. Roughly in Central Asia, R-M207 split into two sister branches one, R1-M173, spread to western Eurasia, while R2-M479, extending southwards, could have entered India through the western or the eastern corridors. The fact that R2 is most concentrated in southern and eastern regions of India^129^ slightly favors the eastern alternative. Finally, the case of H-L901 is remarkably interesting because this basal haplogroup is the only one that seems to have had its first expansion into India. We have already seen that Y-chromosome macro-haplogroup F had its first radiation in Southeast Asia. Afterward, two consecutive splits occurred giving place to the born of haplogroups G and H (SFig 7). Haplogroup G developed a clear western Eurasian phylogeographic pattern with most probable expansion centers situated in the Caucasus or western Iran^132,133^. Curiously, haplogroup G has not been detected in eastern Eurasia and its sporadic presence in India resulted from recent western migrations^128^. As for haplogroup H, it is a specific Indian clade with secondary expansions to eastern and western regions. Although the highest frequencies for haplogroup H were found in southern India^129^, the basal H-M69 haplogroup displayed the highest STR variance in northeast India^129^ which points to an entry through the eastern Indian corridor for the ancestors of Y-chromosome haplogroup H. Human migration into India had to be a complex process as can be deduced by the ample temporal ranges found for the foundation and expansion ages of different mtDNA (Table 1) and Y-chromosome lineages (Table 2). Later Neolithic and post-Neolithic exogenous spreads into India are clearly signaled by Y-chromosome of western (J) and eastern (O) adscription^134^. In addition, it has been observed that these genetic influxes were mediated mostly by males, causing an important sex-bias on the affected populations^135–137^. All these later demic movements have blurred the tracks of the first migrations, but even so, the framework indicated by the uniparental markers is in clear contradiction with the southern route. Furthermore, our genetic hypothesis on the modern human colonization of India finds a better fit into the model deduced from archaeological data, which also question the southern route dispersal of modern humans from Africa to Southeast Asia through India^138^.

### The first colonization of Europe

Archeological findings attest that the first forays of modern humans into Europe took place more than 50 kya^45^. The subsequent Paleolithic colonization movements are well documented by the European archaeological and anthropological records. Recently, notable improvements in the extraction and analysis of DNA from ancient remains has made possible genetic studies on the same samples characterized and dated previously by the anthropologists up to ages close to 50 kya^139^. Thus, in the case of Europe, the human genetic prehistory of their uniparental markers can be directly approached from existing samples along the different archaeological horizons instead of inferring them from the phylogeny and phylogeography of the uniparental lineages present in its current population.

The most ancient European human specimens from Early Upper Paleolithic belonged to eastern Europe and harbor mtDNA macrohaplogroup N basal lineages without present-day direct descendants^139^, and the first recognizable derived N lineage appeared in Crimea as a pre-N1b lineage in a Proto-Gravettian substrate^140^. Mature N1a and N1b lineages as well as X2 and branches I and W1, respectively derived from N1 and N2 trunks, first appeared in the Middle East in Mesolithic times (Table 1). As the 34 ky old Salkhit specimen from Mongolia also harbors a N basal lineage^54^, that had common phylogenetic roots with Bulgarian Paleolithic specimens (SFig. 2: N*3 tree), the most parsimonious hypothesis is to suppose that a basal mtDNA N lineage arrived at Europe, and later to the Middle East along the Caucasus^141^, from a central Asian ancestral population. The Y-chromosomes that potentially accompanied this female westward migration were basal F* and I* lineages (SFig. 7).

Mitochondrial DNA macro-haplogroup M lineages detected in western Asia current populations are derived lineages whose roots are in eastern Asian regions^83^. Surprisingly, basal M lineages were extracted from Early Upper Paleolithic remains in eastern^50^, and Western Europe^142^. They were also found in southern^143^ and southwestern^52^ Mediterranean areas along with Gravettian lithic artefacts. However, after the LGM, M lineages were only detected in Pleistocene northern African contexts as M1b derivatives^144^. In contrast to the case of mtDNA macro-haplogroup N, the Y-chromosome counterparts of these M maternal lineages were in majority C1a lineages (SFig. 7), which points to the possibility that this migratory wave could have followed a more southern route than that used by macro-haplogroup N female carriers, although the coalescence ages of both pioneer groups overlapped.

Some females carrying basal R* lineages could have accompanied to the N* migrants as basal R* types were detected in Early Paleolithic Russian sites^55^, and in eastern^50^ and southern European^145^ Paleolithic contexts. The coexistence of these female lineages with Y-chromosome F* basal lineages (Fig. 2) is favoring again a Central Asian origin for these incomers. Although, perhaps, derived “in situ” from these ancestral R* lineages, the subsequent radiation of mtDNA R* branches until Mesolithic times offer a singular perspective of the regional interactions that occurred in Europe before the Neolithic influences. The most prominent of these R* derived lineages was mtDNA haplogroup U*. Basal U* lineages have been found since the proto-Gravettian in Eastern Europe^146^ and later in Siberia^147^, which again points to an equidistant center of radiation in Central Asia. MtDNA haplogroup U* split into three main independent clusters: U5, U6 and U2’3’4’7’8’9. Nowadays U6 is predominantly found in northern Africa and the European Mediterranean area^148^, but ancestral U6* lineages have been detected in Paleolithic Eastern Europe^149^ and later in Upper Paleolithic Georgia^150^ sites that preceded the LGM. After this drastic period U6 disappeared from Europe but it was persistently detected in Pleistocene remains from Morocco^144^ as derived U6a7 types still present today in the region. These data favor an entrance into northern Africa of U6 following a northern route across the southern Caucasus. The evolution and dispersals of haplogroup U5 along Europe seems to have been overly complex. The earliest U5* basal lineages in Europe have been detected mainly in Central and Western Europe at Gravettian sites, accompanied by F* and C1a2 male lineages^52,143^. These lineages persisted after the LGM, as undefined U5* types until Magdalenian times in Italy^143^ and, most probably, matured in Europe giving place to the two present day main branches U5a and U5b. The U5a branch radiated in Mesolithic times mainly from Eastern Europe, reaching northern Europe at that time^151^. As for the other branch, U5b, basal lineages have been detected in the Iberian Peninsula associated with Magdalenian culture^143^. More derivate, U5b1 lineages were found mainly in Western and Central Europe also in a Magdalenian context^52^, while the U5b2 branch first appeared in the Epigravettian of Italy^142^. This cluster had a generalized expansion at the Mesolithic being present in Europe from west to east. Y-chromosome I2 and R1b lineages were the most representative male counterparts of these female spreads (SFig. 7). As to the U2’3’4’7’8’9 composite branch, its oldest detection occurred at the far East of Siberia, being accompanied by P1-M45 Y-chromosome lineages that were the precursors of eastern and western Asian lineages Q-M242 and R-M207 respectively^152^. In Europe, this undifferentiated lineage is first detected in

Mediterranean regions^52,142,153^, coming accompanied, in addition to the aforementioned I2, by derived Y-chromosomes of the prominent Eastern Asian clade C-M130. Curiously, mtDNA haplogroups U2* and U8* lineages arrived at Europe before than its precursor U2’3’4’7’8’9. They were both present in Eastern Europe since the Initial Upper Paleolithic, being paired with male lineages C1a and C1b of Eastern Asia origin^50,143^. Other lineages reached or expanded in Europe during the Mesolithic as mtDNA haplogroups H7, H13, K, R1b, or U4, and Y-chromosome haplogroups I2a1, I2a2, J1, R1a or R1b (SFig. 7). Finally, other lineages considered of Neolithic adscription in Europe were present in the Middle east at least since the Mesolithic as is the case of mtDNA lineages H*, H5, HV, HV2, I, J, K, N1a, R0a, T, U3, U7, W, or X2 and Y-chromosome haplogroups G2a, G2b, J1, J2, or T.

In short, and as previously stated^85,154^, Europe, the westernmost Peninsula of Asia, was occupied by modern humans later than the rest of the Continent, and its colonizers probably reached the region through the Eurasian Steppe first, and through the Near East latter, which in turn was presumably colonized by a parallel southern wave from Central Asia that reached the region through Iran and bordering the Caucasus.

## Conclusions

Phylogenetic and phylogeographic analyses of uniparental genetic markers on present and past human populations, under the perspective of an evolutionary rate slowdown going back in time, allowed the construction of demographic models that explain the first spread of modern humans across Eurasia, Australasia and Near Oceania in harmony with the archaeological and fossil records.

